# Bioengineering of Pea (*Pisum sativum*) for the Expression of Myoglobin, a Heme-containing Animal Protein

**DOI:** 10.64898/2026.09.01.748490

**Authors:** Rishikesh Ghogare, Bruce Williamson, Karen Adams, Nathan Tarlyn, Kamryn Clements, Daniel Velazquez, Lydia Wilcox, Raechel Holbrook, Justin Helm, Nathaniel Gruca, Amit Dhingra

## Abstract

Myoglobin, an oxygen-binding animal protein, was engineered in *Pisum sativum* (pea) to explore its potential as a food ingredient and balance the amino acid profile. In this study, minimal expression cassettes and binary vectors were used to express bovine myoglobin using particle gun and Agrobacterium-mediated transformation, respectively. Successful integration and expression of the myoglobin gene was achieved in *P. sativum,* with both methods yielding similar transformation efficiencies (∼1%). Expression analysis of T2 seeds revealed that Agrobacterium-mediated transformation-derived transgenic lines that expressed myoglobin under the regulation of a Soybean 7S seed-specific promoter and Tobacco Etch Virus (TEV) translation enhancer and a chimeric Rb7MAR Terminator (Ps-BpRG13 events) consistently yielded the highest level of expression (0.32–1.57% of TSP), while transgenic lines with myoglobin expression under the regulation of a Soybean Phaseolin promoter and Rb7MAR Terminator (Ps-BpRG14 events) resulted in moderate levels of heterologous protein expression (0.13–0.83% TSP). Transgenic events with constitutive 2xCaMV35S promoter, TEV translation enhancer and Rb7MAR terminator (Ps-BpRG15 events) exhibited the lowest level of myoglobin expression (0.09–0.14% TSP). Co-bombardment of two minimal expression cassettes – one with myoglobin under the regulation of the Phaseolin promoter and Rb7MAR Terminator and the other with the nptII selectable marker under the regulation of a 2X constitutive CaMV35S promoter, TEV translational enhancer and TNOS Terminator, yielded lines that exhibited variable expression (0.03–0.77% TSP), with some events comparable in expression to Agrobacterium-derived Ps-pRG14 events. To the best of our knowledge, this is the first report of producing a heme-containing animal protein, myoglobin, in peas, with potential implications for sustainable production of food ingredients and nutritionally fortified and value-added plant products using molecular farming.

## Introduction

It is estimated that by 2050, 80% more protein will need to be produced to meet the demand of an increasing human population (Henchion et al. 2017). Currently, over 65% of the protein supply in the human diet is derived from animals with the remainder from plants. Recently, plant-derived proteins have increasingly become popular for ethical, environmental, and health reasons. Most of the plant-derived proteins are sourced from Soybean (*Glycine max*) and Pea (*Pisum sativum*). However, plant-derived proteins have nutritional limitations compared to animal-sourced proteins (Day et al. 2022; Soh et al. 2025). Some of the key challenges associated with plant-based proteins in human nutrition include incomplete essential amino acid profile, digestibility, and bioavailability (Partanen et al. 2025). Bioengineered crops that produce selected animal proteins and satisfy the nutritional, organoleptic, and functional traits that consumers prefer can help in meeting the global protein demand (Messina and Messina 2024).

With the aim of their use as a food ingredient, several animal proteins have been successfully expressed in different tissues of a wide range of plant species in the academic and private sectors (Tusé et al. 2024). The plant-expressed proteins include lactoferrin, various caseins present in milk, and chymosin. These have been expressed in the seeds of rice, soybean, safflower, and lettuce leaves (Philip et al. 2001; Nandi et al. 2002). The egg white protein, ovalbumin and several structural muscle proteins have also been successfully expressed in bioengineered crops (Wandelt et al. 1991). Collagen and myosin have been engineered in the leaves of lettuce, kale, and spinach, and the muscle oxygen-carrier myoglobin in corn and soybean seeds (Shoseyov et al. 2013; Tusé et al. 2024).

Of all these proteins, Myoglobin (Mb) plays a major role in the texture, appearance, and flavor of the animal meat during consumption. Mb is responsible for the red color in meat and when exposed to oxygen it gives fresh meat a bright red appearance which turns brown when meat is cooked due to oxidation (Suman and Joseph 2013). It also contributes to the iron-rich “meaty” flavor of traditional meat and overall juiciness and appeal. In addition, Mb is rich in bioavailable iron, which is one reason meat is a significant source of iron in the human diet (Devaere et al. 2022). Therefore, myoglobin or a plant-based equivalent is considered important in food formulations (Fraeye et al. 2020). In a recent study, Mb was expressed in the cytosol and chloroplast of plant cells using various expression systems, including viral vectors and gene fusion constructs (Carlsson et al. 2020). The expressed myoglobin was functional in terms of ligand binding, indicating that plants can support the expression of functional myoglobin. Successful demonstration of myoglobin expression in plants unlocks the potential for its utilization in the food industry and beyond. Soybean has been reported to express porcine myoglobin protein at over 25% TSP (Paladini et al. 2024). There are advantages to expressing myoglobin in pea as it represents the number one plant protein in new vegan products with a projected compound annual growth rate of 12.78%, and importantly allergenicity issues common with Soybean (Davis, 2025).

Belonging to the Fabaceae family, Garden Pea, *Pisum sativum*, has been used due to its ease of cultivation and short lifecycle (Ludvíková and Griga 2022; Yang et al. 2022). Peas are considered an important source of protein, starch and fiber. Due to relatively high protein content and protein properties such as low allergenicity, high water solubility, oil holding capacity, emulsion ability and gelation, they are potential ingredients in the processed food industry. In addition to processed foods, pea protein can also be used as emulsifier, encapsulating material and biodegradable natural polymer (Shanthakumar et al. 2022) . The recent reports related to efficient Agrobacterium-mediated transformation and CRISPR-Cas9 mediated genome editing has opened up possibilities to express animal proteins and for the development of varieties with better agronomic traits (Kaur et al., 2022; Li et al., 2023; Williamson-Benavides et al., 2025).

While transformation is now feasible, expression of complex heme-containing animal proteins in pea seeds remains underexplored. In this study, we expressed *Bos taurus* (Bovine) Mb gene in *Pisum sativum* using both particle bombardment and agrobacterium-mediated transformation under the control of constitutive and Soybean seed specific promoters. To the best of our knowledge, this is the first report of expressing Mb in pea as well as demonstrating successful particle gun mediated transformation using minimal transgenic expression cassettes. The two bioengineering approaches resulted in a total of 85 independent T0 transgenic events. In the T2 generation, some events resulted in myoglobin accumulation exceeding 1% TSP reaching 1.57% TSP in the seeds of one of the transgenic lines.

## Materials and Methods

### Synthesis and cloning of Mb gene and construction of plant transformation vectors

The *Bos taurus* (bovine) Mb gene sequence was acquired from the Uniprot database (accession number NP_776306) and codon optimized for expression in pea using SnapGene V.6 (San Diego, CA). Codon-optimized Mb gene and legume-specific β-Phaseolin (PPHAS) promoter, 7S globulin promoter fused with the Tobacco Etch Virus translational enhancer (PPHAS and P7STEV), and a customized chimeric ARC5 terminator attached with tobacco Rb7 matrix attachment region (Rb7MART) were synthesized at Genscript (Piscataway, NJ, USA). The seed-specific promoters 7S and PHAS were derived from genes encoding 7S globulin and β-Phaseolin from *Phaseolus vulgaris* and *Glycine Max*, respectively (Zakharov et al. 2004; Fauteux and Strömvik 2009). The DNA sequence of the codon-optimized Mb gene is available in the supplementary material and the DNA sequence for the regulatory elements is the same as reported previously (Paladini et al. 2024). All recombinant DNA manipulations were performed in *Escherichia coli* strain Top10. Sequences of all oligonucleotides used in the study are listed in **Table 1**.

**Table 1.**
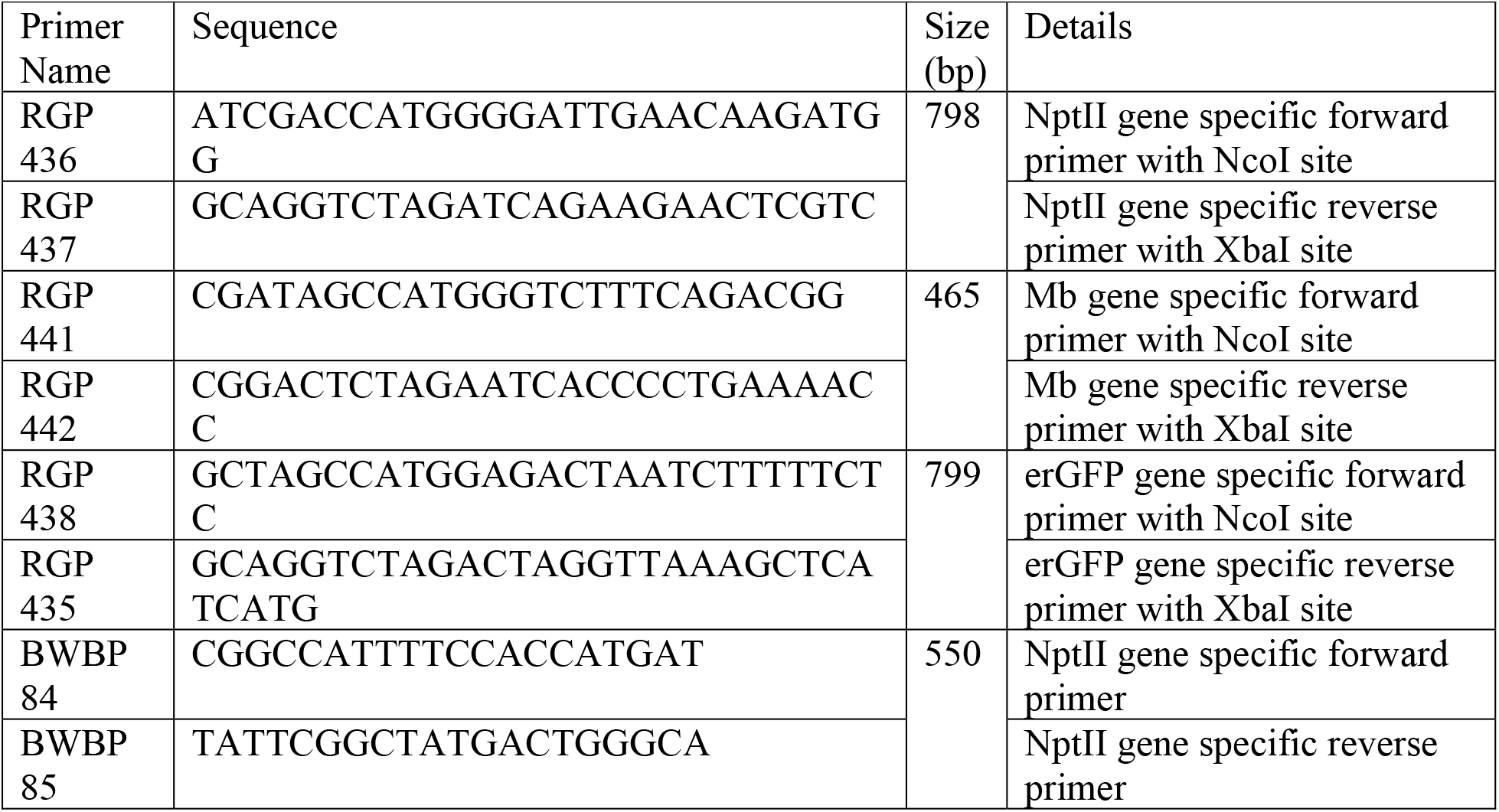
List of oligonucleotides used in the study.

The pUC57 vector containing the synthetic Mb gene, P7S-TEV promoter, and Rb7MarT terminator (pRG13) (**Figure 1**) was synthesized by Genscript (Piscataway, NJ, USA) with the sequence immediately upstream of the translation initiator ATG codon modified to an *NcoI* site, while the stop codon altered to an *XbaI* restriction site. The Mb gene was released from pRG13 using *NcoI/XbaI* and cloned into synthesized pUC57 vector containing PPHAS promoter and Rb7MAR terminator in *NcoI/XbaI* site between the promoter and terminator (pRG14) (Figure 1). The Mb gene digested with *NcoI/XbaI* was used to replace smGFP in pAD120 derived from pAVA121 containing a double CaMV35S promoter, Tobacco Etch virus translational enhancer and Nopaline synthase (NOS) terminator (pRG15) (**Figure 1**) (Jiwan et al. 2013).

**Figure 1.**
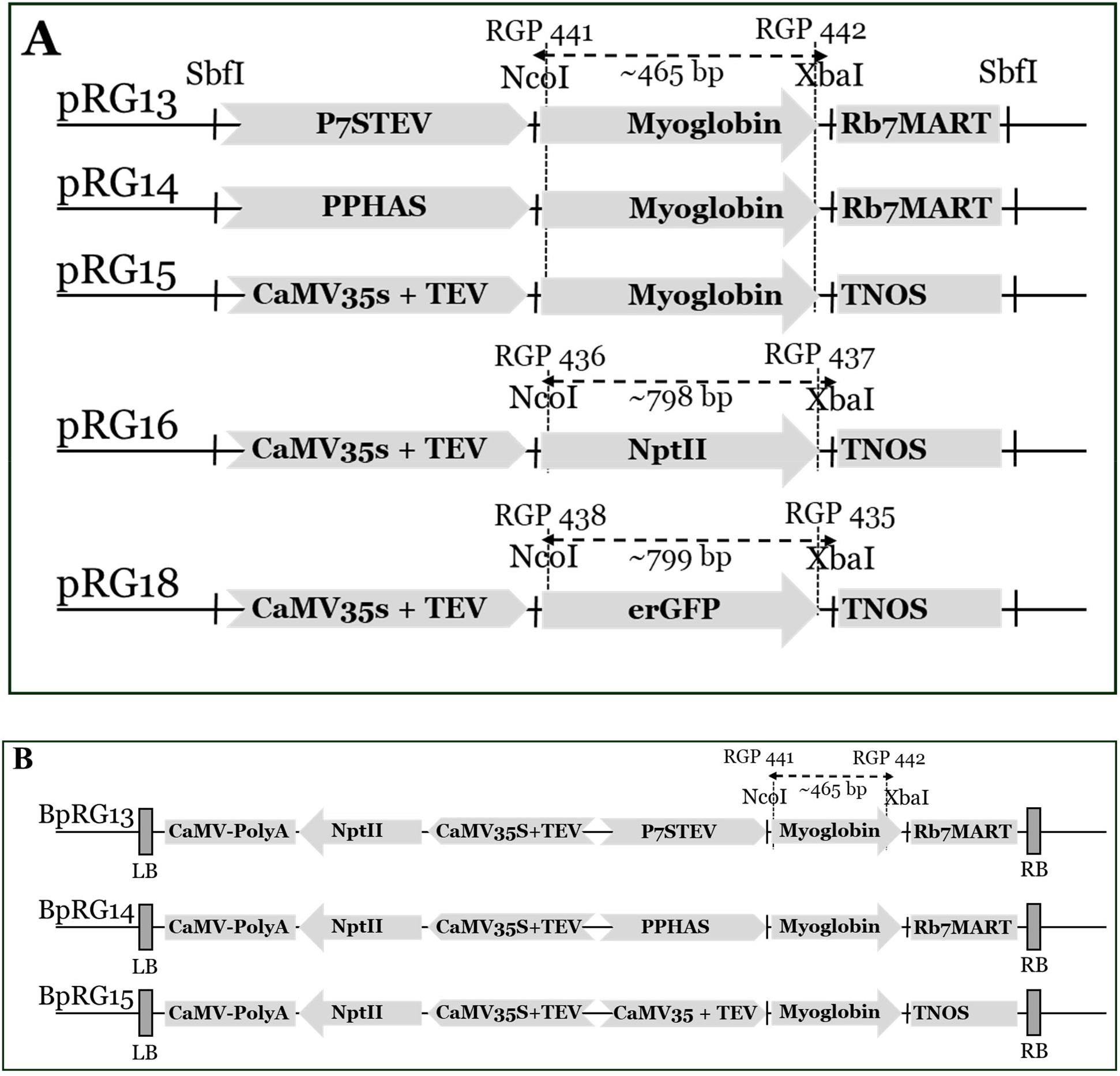
Schematic representation of plant transformation vectors used in this study. A. Plasmids used in biolistic transformation; B. Binary vectors used for agrobacterium-mediated transformation. Abbreviations: Mb, myoglobin; NptII, neomycin phosphotransferase II (kanamycin-resistance selectable marker); P7S-TEV, soybean 7S globulin seed-specific promoter fused to the Tobacco Etch Virus (TEV) translational enhancer; PPHAS, Phaseolin seed-specific promoter; CaMV35S, Cauliflower Mosaic Virus 35S constitutive promoter; Rb7MART, chimeric ARC5 terminator fused with the tobacco Rb7 matrix attachment region; NOS, nopaline synthase terminator.

For use in particle gun mediated transformation, minimal selection cassettes needed to be prepared. For this step, the Kanamycin resistance (NptII) gene was amplified from plasmid pCambia 2300 using primer RGP 436 and RGP 437 (**Table 1**). PCR was performed using the NEB Q5 high fidelity DNA polymerase. The primers RGP 436 and RGP 437 were used to introduce a *NcoI* site at the 5’ end of the gene and an XbaI site at the 3’ end of the gene, respectively. The NptII gene amplicon was digested with *NcoI/XbaI* and cloned into pAD120 at the *NcoI/XbaI* site as described before (pRG16). Similarly, as a control, erGFP gene from plasmid CH2-185 was amplified using primers RGP 435 and RGP 438 and cloned into pAD120 digested with *NcoI/XbaI* to develop plasmid pRG18 (**Figure 1**).

To develop binary transformation vectors, the entire expression cassette including promoter-Mb gene – terminator was released from pRG13, pRG14, and pRG15 using *SbfI* and cloned in *SbfI*-digested pCambia 2300. The resulting plasmids are referred to as BpRG13, BpRG14 and BpRG15, respectively with B denoting the binary pCambia vector (Figure 1). All the resulting binary plasmids were confirmed by restriction digestion and full plasmid DNA sequencing at MGH DNA Core, MA. The plasmids were then introduced into Agrobacterium strain GV3850 using electroporation.

### Plant material and Explant preparation

Production of the plant material and explant preparation were performed as described earlier (Williamson-Benavides et al. 2025). Briefly, *Pisum sativum seeds* (accession no.-PI 175226) were obtained from the USDA Western Regional Plant Introduction Station (Pullman, WA, USA). The seedlings were germinated in 2.8 L pots containing Sunshine Mix#1/LC1 (Sun Gro Horticulture, Agawam, MA, USA). Plants were maintained in a climate-controlled greenhouse at Washington State University Plant Growth Facilities with temperature held at 24 °C / 18 °C (day/night) and relative humidity maintained at 40-60%. The day length conditions were 16h day and 8 h night. Plants were grown until immature pods were formed and then pods were harvested. The immature pod stage is also known as “eating pea” stage, at which seeds have reached maximum size but have not started to dry. Immature pods (9-12 days old) were collected and stored at 4 °C for a period no longer than a week. The pods were then incubated in 70% (v/v) ethanol for 2 min with constant agitation. This was followed by rinsing once with distilled water, then immersion in a fresh 1% (w/v) sodium hypochlorite solution with 1-2 drops of Triton X-100 for 20 min. Surface sterilization was completed with three washes of distilled water. Seeds were then recovered from the pods and excised along the embryonic axis, separating the cotyledons. The root end of the embryo and 80% of the cotyledon distal to the embryonic axis were excised and discarded. Then the excised embryo was then placed on pea regeneration media (PRM: Murashige and Skoog (MS) with vitamins, 3% sucrose, 0.2% Gelrite, 5 mg/L Indole-3-butyric acid (IBA),0.5 mg/L 6-Benzylaminopurine (BAP) and pH 5.8) for 8 days before particle bombardment or co-cultivation with Agrobacterium. Shoots developed from the embryo during the pre-treatment period were removed before the transformation procedure.

### Agrobacterium-mediated transformation

*Agrobacterium* strain GV3850 harboring either BpRG13 or BpRG14 or BpRG15 or pCambia2300 (empty vector) was used to inoculate 2mL LB media supplemented with 100 mg/L Kanamycin and incubated overnight at 28 °C. The starter culture was used to inoculate 200 mL LB containing 100 mg/L Kanamycin and incubated at 28 °C for 48h. The *Agrobacterium* cultures were then centrifuged at 5000 rpm for 10 min and the pellet was re-suspended in liquid co-cultivation media (MS w/ vitamins, 100 g/L sucrose, 5mg/L IBA, 0.5 mg/L BAP, 200μM acetosyringone and pH 5.8) until the OD_600_ reached 1.6.

The pre-treated explants were transferred to the Agrobacterium culture and incubated for 1h at 28°C with gentle agitation. After co-cultivation, the explants were blotted on sterile filter paper to remove excess agrobacterium suspension and transferred to solid co-cultivation media (MS w/ vitamins, 30 g/L Sucrose, 0.2% Gelrite, 5mg/L IBA, 0.5 mg/L BAP, 200 µM acetosyringone and pH 5.8). The explants were incubated in the dark at 24°C for 48h. After co-cultivation the explants were rinsed three times with sterile water supplemented with 250 mg/L Timentin. Explants were then incubated in sterile water supplemented with 400 mg/L Timentin for 1h at 28°C with gentle agitation. After washing, the explants were blotted on sterile filter paper and transferred to PRM media supplemented with 250 mg/L Timentin and 20 mg/L Geneticin. The explants were sub-cultured every 2-3 weeks on different formulations of selection media. The initial round of selection used PRM media supplemented with 250 mg/L Timentin and 20 mg/L Geneticin and the second round of selection used PRM media supplemented with 250 mg/L Timentin and 25 mg/L Geneticin. The third and fourth rounds of selection used PRM media supplemented with 250 mg/L Timentin and 30 mg/L Geneticin. Next, regenerated shoots were transferred to pea elongation media (PEM: ½ MS w/ vitamins, 30 g/L sucrose, 0.2% gelrite, 0.1 mg/L Indole-3-acetic acid (IAA), 300 mg/L Timentin and 30 mg/L Geneticin) for two rounds of subculture. After elongation the shoots were transferred to rooting media (PROOT: MS w/ vitamins, 16 g/L glucose, 0.2 % Gelrite and 100 mg/L Timentin) until rooted plants were obtained. Agrobacterium-mediated transformation was performed across 7–9 independent experiments per plasmid, using a larger number of explants (an average of 4,284 explants per plasmid). Explants co-cultivated with control empty vector were processed similar to the explants treated with myoglobin expressing plant transformation vectors.

### Particle Gun-mediated Transformation

The plasmids pRG14, pRG18 and pRG16 were digested with *SbfI* to release the minimal expression cassette containing promoter-gene-terminator for the Mb gene and NptII gene, respectively. The released minimal expression cassettes were purified and concentrated using gel purification. The individual minimal expression cassettes for the Mb gene and NptII gene were then re-circularized using T4 DNA ligase before coating the gold particles. The gold particles suspension was prepared as reported previously (Dhingra and Daniell 2006).

The circularized minimal expression cassettes for the Mb gene (1.5 µg) and NptII gene (1.5 µg) were mixed in equimolar ratio and used for coating 0.6µm gold particles. Briefly, the circularized expression cassette was added to 50 µL gold suspension. Then, a total of 50μL sterilized 2.5 M CaCl_2_ and 20μL 0.1M spermidine was added to the DNA-gold solution and the mixture was vortexed for 20 min at 4°C. An aliquot of 200μL 100% ethanol was added to the mixture, vortexed briefly, and centrifuged at 3000 rpm for 30 sec. The supernatant was discarded, and the pellet was washed three more times using 100% ethanol. After the final wash, the pellet was re-suspended in 35 µL of 100% ethanol. A total of 10μL DNA-gold mixture was used for each bombardment (Dhingra and Daniell 2006). Similarly, circularized minimal expression cassettes for GFP gene (1.5µg) and NptII gene (1.5µg) were also applied to gold particles as a bombardment control. The biolistic bombardment was performed using a PDS1000/He particle bombardment system (Bio-Rad) with a target distance of 6.0 cm from the stopping plate at a helium pressure of 1,200 psi. After bombardment, explants were placed in the dark for 48h at 25 °C for recovery. Finally, the explants were transferred to selection media as detailed previously (Williamson-Benavides et al. 2025). For each minimal expression cassette, gene gun–mediated transformation was carried out across 2–3 independent experiments, using 315 to 1,139 explants per plasmid in total. Each experiment involved a discrete batch of explants subjected to biolistic delivery of the corresponding construct.

### Identification of transgenic events

The tissue from young leaves was collected for DNA extraction. Total cellular DNA was extracted using the DNeasy Plant Mini Kit (Qiagen, Hilden, Germany). Gene-specific primers RGP 441 and RGP 442 **(Table 1)** for the Mb gene and RGP 436 and RGP 437 for the NptII gene (**Table 1**) were used for the PCR assay. The PCR conditions were set as follows: one cycle at 95°C for 3 min, followed by 35 cycles of 95°C for 30 s, 53°C for 30 s, 72°C for 1 min, and a final extension at 72°C for 5 min. DreamTaq DNA polymerase (Thermo Fisher, US) was used for the PCRs. PCR was used only to confirm transgene presence and not to determine copy number, zygosity, or site of integration.

### Quantification of myoglobin expression

The transgenic T0 plants transformed with pCambia2300, pCambia 2300-pRG13, pCambia 2300-pRG14, pCambia 2300-pRG15, and pRG14+16 constructs were cultivated and propagated to T2 seeds. T2 seeds were screened for the presence of the NptII gene via PCR using primers BWBP 84 and BWBP 85 (**Table 1**). Total cellular DNA was isolated using the CTAB method from a small section (25 mg) of the seed.

Prior to extraction, dry T2 seeds were soaked in MilliQ water for 16 h to imbibe the tissue, and 50 mg of the resulting hydrated tissue was used per sample. T2 Transgenic seeds were processed in protein extraction buffer (5% w/v SDS, 175 mM Tris-HCl, pH 8.0, 0.4% v/v beta-mercaptoethanol) with Omni ceramic beads (1.4 mm). The extract was incubated at 65 °C for 25 min, centrifuged and then the supernatants were transferred to fresh tubes. The extract was diluted 1:25 in water and total protein concentration was determined using the Pierce™ Detergent Compatible Bradford Assay Kit (Thermo Fisher Scientific, Cat# 23246) according to the manufacturer’s protocol.

Myoglobin quantitation from seed extracts was done using the Bovine MB / Myoglobin (Sandwich ELISA) ELISA kit (LSBio, Cat# LS-F51253-1). All samples were normalized to 50µg/mL Total Soluble Protein (TSP) myoglobin content was measured according to the manufacturer’s protocols. The concentration of myoglobin was determined by reference to the standard curve. Each event consisted of at least one biological replicate with three technical replicates. 20 µL (1 µg) of sample was used for each test. Biological replicates corresponded to independent T2 seed pools from individual events, while technical replicates represented repeated ELISA measurements from the same extract. Heme incorporation or oxygen-binding functionality of myoglobin was not assessed in this study.

### Statistical Analysis

Three technical replicates for each biological replicate or transgenic event were used to quantify myoglobin expression with ELISA. Since independent biological replication at the event level was not feasible and thus not performed, no formal inferential statistical test was applied to compare myoglobin accumulation among constructs, regulatory elements, or transformation methods. Table 4 presents descriptive data as the mean ± standard deviation (s.d.) of the three technical replicates for each event. The number of independent transgenic events assayed per construct is reported in Table 4, and the number of explants screened, and transgenic events recovered per transformation method and per experiment are reported in Tables 2 and 3 and in the Materials and Methods.

**Table 2.** Genetic transformation of *Pisum Sativum* using particle bombardment method. Verified transgenic events were confirmed by PCR for the Mb (or NptII, for the pRG18+16 GFP/marker control) transgene using primers listed in Table 1 (see Materials and Methods, Identification of transgenic events); PCR confirmed transgene presence or absence. Copy numbers or zygosity were not determined. Data represent totals across 2–3 independent bombardment experiments per plasmid combination.

| Plasmid (co-bombardment) | Number of explants | Number of verified transgenic events | Transformation efficiency (%) |
| --- | --- | --- | --- |
| pRG14+16 | 487 | 5 | 1.02 |
| pRG18+16 | 159 | 0 | 0 |

Transformation efficiency = (Number of verified events/Total number of explants) \*100

**Table 3.** Genetic transformation of Pisum Sativum using *Agrobacterium.* Verified transgenic events were confirmed by PCR for the Mb transgene using primers listed in Table 1 (see Materials and Methods, Identification of transgenic events); PCR confirmed transgene presence only, not copy number or zygosity. pCambia2300 denotes the empty-vector (no myoglobin cassette) negative control. Data represent totals across 7–9 independent Agrobacterium-mediated transformation experiments per plasmid (see Materials and Methods, Agrobacterium-mediated transformation).

| Plant Transformation Vector | Number of explants | Number of verified transgenics | Transformation efficiency (%) |
| --- | --- | --- | --- |
| BpRG13 | 3024 | 19 | 0.63 |
| BpRG14 | 3060 | 22 | 0.72 |
| BpRG15 | 3144 | 22 | 0.70 |
| pCambia2300 | 720 | 15 | 2.08 |

Transformation efficiency = (Number of PCR-verified events/Total number of explants) \*100

**Table 4.**
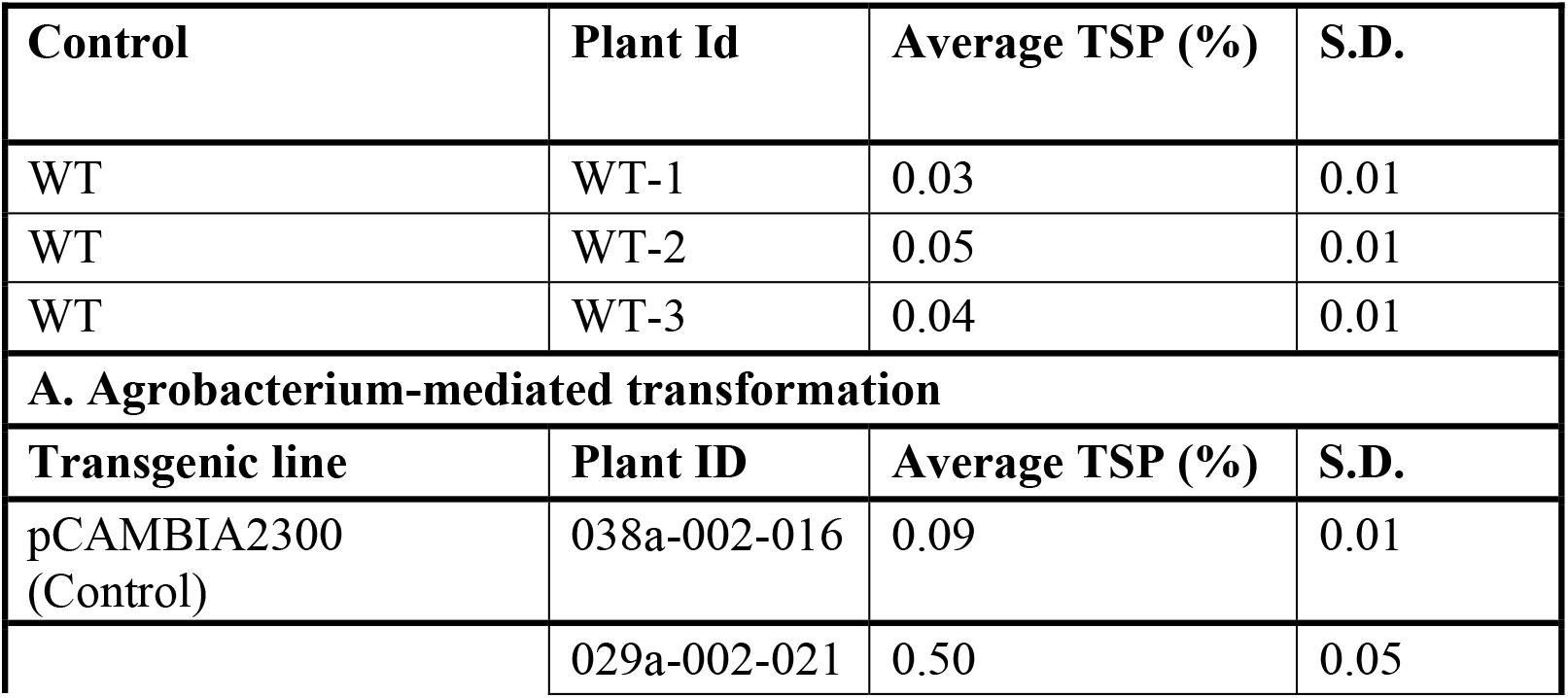
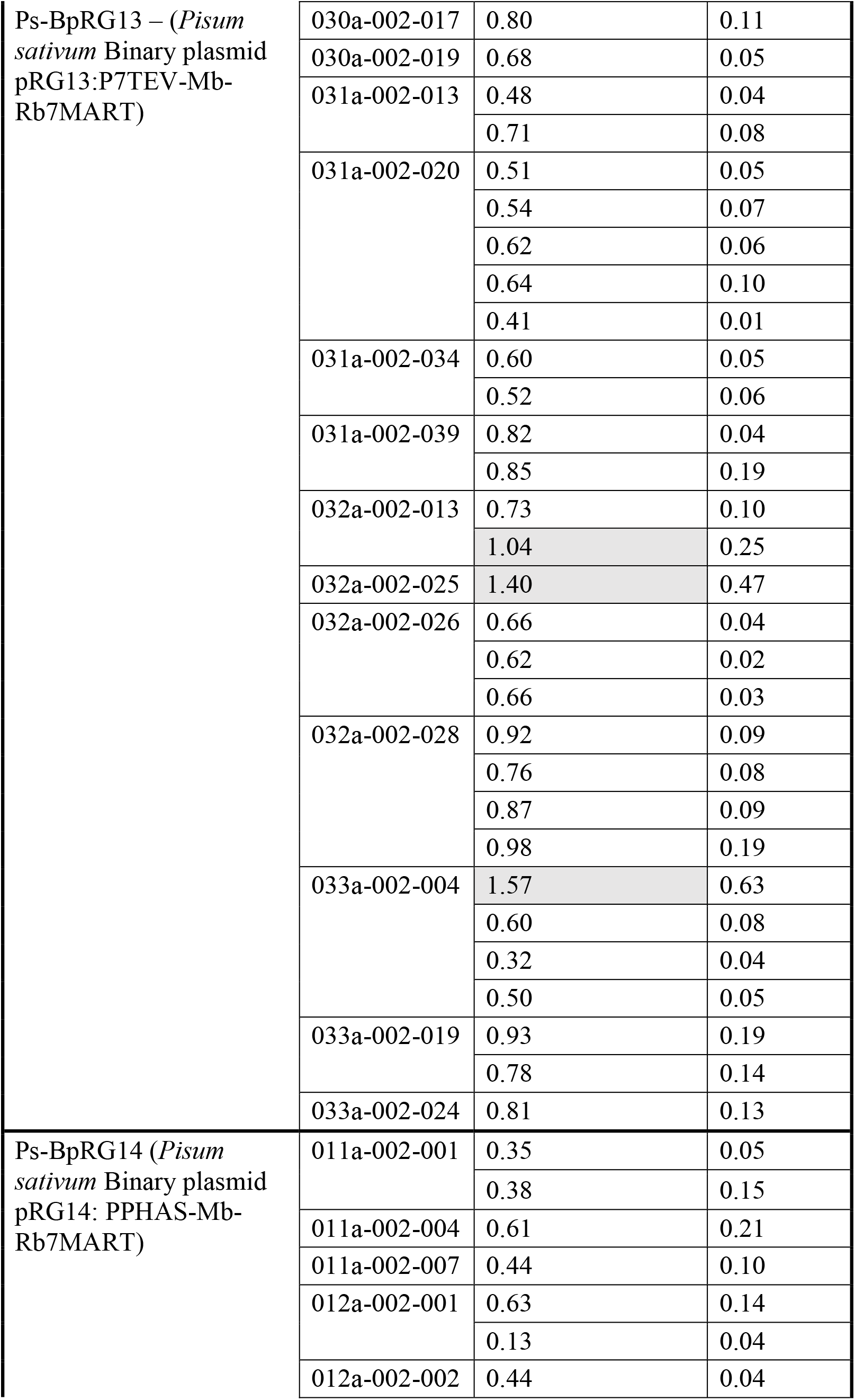
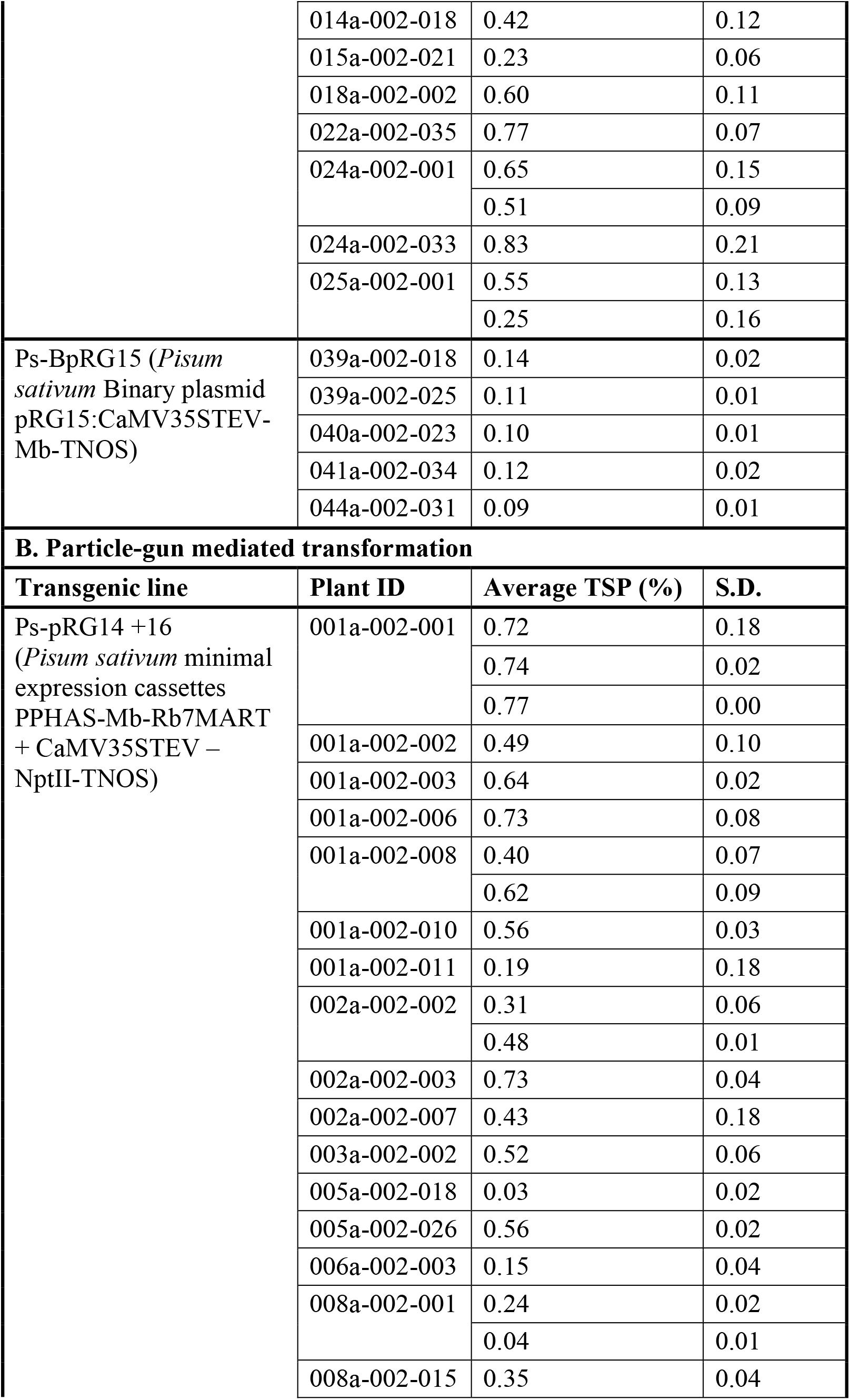
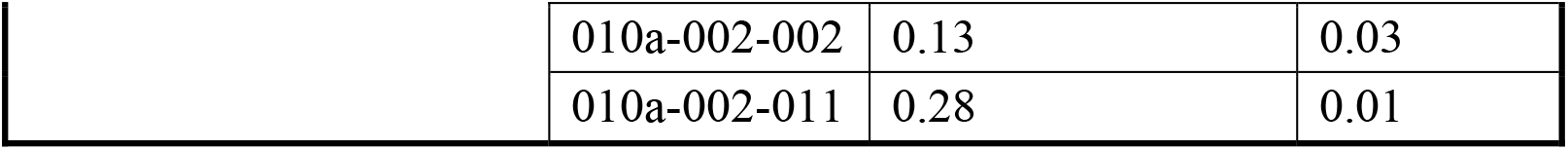
Quantification of myoglobin as a percentage of TSP content (±s.d.) in independent transgenic T2 pea seeds. A. Agrobacterium-mediated transformation, B. Particle gun-mediated transformation. Lines with myoglobin TSP content above 1% are shaded in gray. TSP, total soluble protein; WT, wild-type (non-transformed) negative control; pCambia2300, empty-vector negative control (no myoglobin expression cassette). Values are the mean ± standard deviation (s.d.) of three technical ELISA replicates from a single biological replicate (one T2 seed pool) per event.

## Results and Discussion

The bovine Mb gene was successfully integrated into the pea genome using both Agrobacterium-mediated and particle gun-mediated transformation. The expression was driven by seed-specific and constitutive promoters.

For particle gun-mediated transformation, two separate minimal expression cassettes containing Mb gene (pRG14) and NptII (pRG16) selection marker were co-bombarded into the explants. The rationale of separating the selectable marker from the gene of interest was that it would be feasible to segregate out the selectable marker in the future generations. This assumes that the integration site for the two expression cassettes would be in non-linked regions of the genome. PCR analysis confirmed the presence of Mb transgene in 5 transgenic events resulting in an efficiency of 1.02% (**Table 2**). However, despite several rounds of selection on Geneticin, the sNptII gene was not detected in transgenic plants. This could be due to the presence of NptII expressing transiently in earlier stages of the selection process. As a control, a set of explants were also co-bombarded with minimal cassettes containing GFP (pRG18) and NptII (pRG16) genes. However, no regenerants were obtained. To the best of our knowledge, this is the first report of obtaining transgenic events in pea using separate minimal expression cassettes representing the gene of interest and a selectable marker.

A previous report using particle gun-mediated transformation in pea reported a low transformation efficiency of 0.4% (Kaur et al. 2022b). We obtained a transformation efficiency of 1.02%, which may be improved with complete circular plasmids. There are reports of using particle gun mediated transformation of other legume species with varying success. In common bean (*Phaseolus vulgaris*), particle gun-mediated transformation was used to co-bombard multiple genes. This study reported an overall transformation frequency around 0.9% and approximately 50% of transgenic events had both plasmids (Aragão et al. 1996). In chickpea (*Cicer arietinum*) particle gun-mediated transformation initially reported very low success rates, and with significant optimization a transformation efficiency of 18% was obtained (Indurker et al. 2007; Kaur et al. 2017). Soybean was the first legume species to be transformed using an electrical discharge particle gun (Christou 1990), and there are several reports targeting various traits (Xu et al. 2022). Minimal expression cassette integration was also demonstrated, which yielded simpler transgene insertions (Vianna et al. 2011; Jackson et al. 2012). It is important to note that the transformation efficiency can be impacted by the gene of interest and the genetic background of the plant species. Therefore, while a transformation efficiency of 1.02% was obtained in this study, it may be further improved by standardizing the cultivar and utilizing pea-specific regulatory elements.

A generalized scheme for agrobacterium mediated transformation for the integration of Mb gene in pea is presented in **Figure 2** and is similar to the one as reported recently (Williamson-Benavides et al. 2025). Using this method a total of 19, 22, and 22 rooted transgenic events corresponding to P7STEV (BpRG-13), PPHAS (BpRG-14) and CaMV35S-TEV (BpRG-15) promoter elements were obtained. A total of 15 events were obtained for the control binary vector pCambia 2300. A transformation efficiency of 0.63-2.08% was obtained with the agrobacterium mediated-transformation method, which is marginally lesser than the 2.9% efficiency observed previously in our research (Williamson-Benavides et al. 2025). Overall, both transformation methods resulted in comparable efficiencies (**Table 3**). While the particle bombardment approach is limited by some parameters such as the need for large DNA quantity, relatively low throughput and limited bombardment area, the use of minimal cassette and absence of NptII gene can potentially save time in developing lines without a selectable marker.

**Figure 2.**
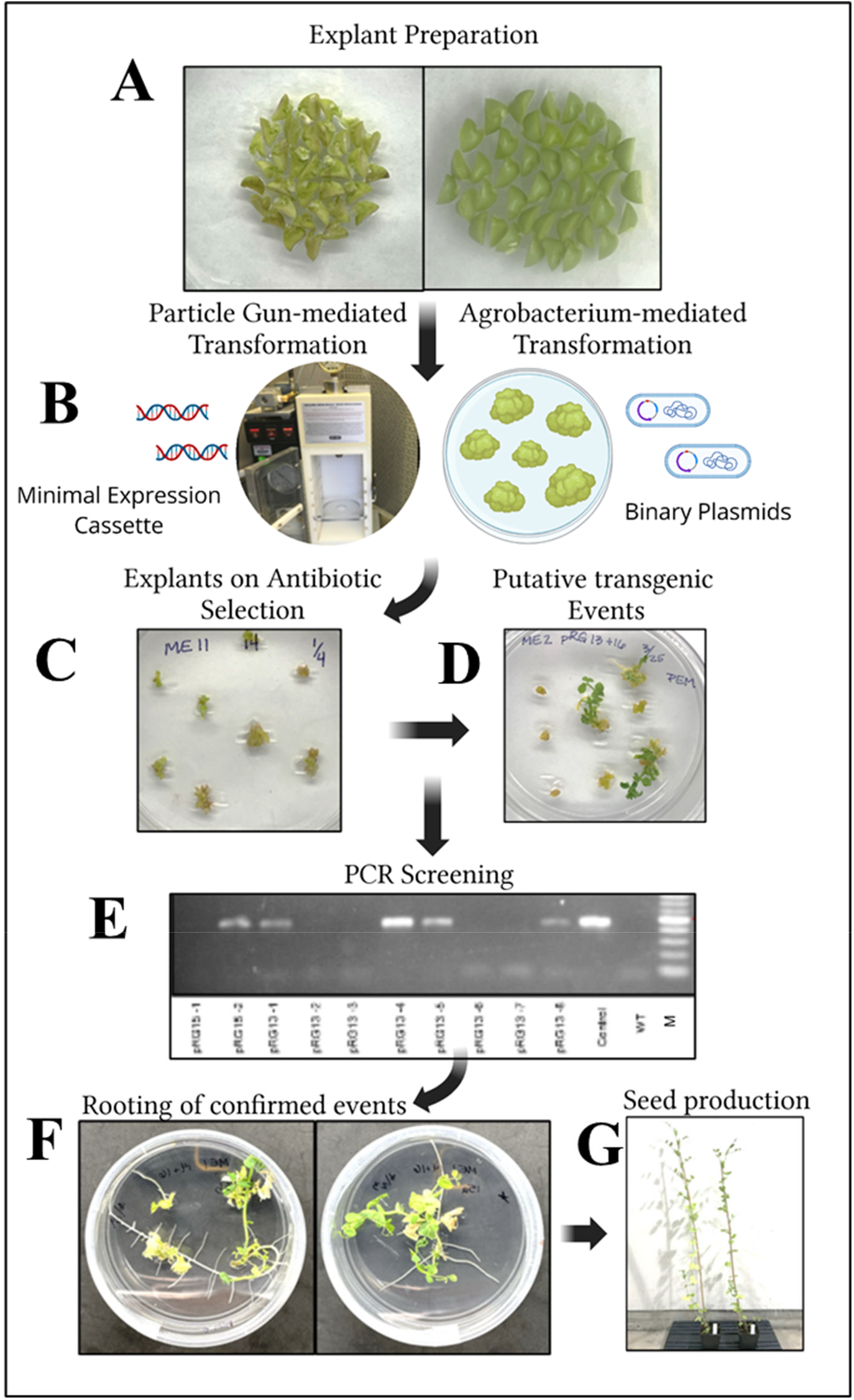
Schematic representation of the plant transformation workflow used in this study to develop pea transgenic lines expressing Mb gene. A. Explant preparation for particle-gun mediated transformation or Agrobacterium-mediated transformation; B. For particle-gun mediated transformation, minimal expression cassettes presented in Figure 1A were used, and for Agrobacterium-mediated transformation, pCambia-based plant transformation vectors presented in Figure 1B were used. C. Explants were placed on selection media (please see materials and methods); D. Putative transgenic events were recovered after a period of selection; E. PCR confirmation of transgenic events using oligonucleotides listed in **Table 1**; F. PCR-confirmed transgenic events were rooted and moved to potting mix; G. Plants were grown out in a growth chamber for collecting seeds.

Agrobacterium-mediated transformation of pea (*Pisum sativum*) has historically shown low efficiency, with reports documenting transformation frequencies ranging from 0.5–2% (Schroeder et al. 1993; Jordan and Hobbs 1993; Grant et al. 1995; Bean et al. 1997). Despite the low efficiency, these studies demonstrated that it is feasible to genetically engineer different cultivars of pea using different types of explants. Protocol refinements that include improved explant selection strategies, hypervirulent Agrobacterium strains, and media additives have modestly increased efficiency to 3% in some cases with one study reporting a 4.1% efficiency (Nadolska-Orczyk and Orczyk 2000; Pniewski and Kapusta 2005).

Pea has been bioengineered to express several foreign genes ranging from ones that provide resistance to insects, viruses and other diseases, abiotic stresses, and for quality and productivity traits. It has also been used as a platform for molecular farming to produce single chain Fv (ScFV) antibody fragment in pea seeds for cancer diagnosis and therapy (Perrin et al. 2000). Another study was able to achieve upto 2% TSP of the ScFV antibody (Saalbach et al. 2001). Follow up studies demonstrated the functionality of pea seed-derived ScFV antibodies (Zimmermann et al. 2009). One of the events under the regulation of 7S globulin promoter yielded 1.57% TSP of myoglobin. This indicates that with screening of additional transgenic lines, or use of pea-specific regulatory elements, such as the USP promoter along with LeB4 signal used by Saalback et al., 2001, it is possible to achieve higher levels of foreign protein accumulation. Especially with the availability of the pea genome, and various seed-related gene expression studies, it is feasible to identify regulatory elements that will enhance the expression of foreign genes in pea seeds (Chen et al. 2013; Liu et al. 2015; Malovichko et al. 2020; Williamson-Benavides et al. 2020). In another study, the VP60 coat protein from the rabbit hemorrhagic disease virus was fused to Cholera Toxin B subunit and expressed in pea. The resulting oral vaccine was tested, and it was shown to elicit immunity in animal tests (Mikschofsky et al. 2009). In another interesting approach, the human acidic fibroblast growth factor (aFGF) was transiently expressed in pea using vacuum-infiltration. The gene was expressed under the regulation of CaMV35S promoter and was introduced using a pea early browning virus-based binary vector that is expected to multiply post-infiltration and amplify the expression (Fan et al. 2011).

### Quantification of myoglobin in T2 Seeds

Expression of myoglobin in transgenic pea seeds varied markedly depending on the construct and transformation method used. Wild-type (WT) controls and empty vector (pCambia2300) lines accumulated negligible levels of myoglobin, averaging 0.03–0.09% of TSP, confirming the absence of endogenous expression and validating the transgenic approach.

Among the transformation constructs, pRG13 (P7STEV Promoter) transformed via Agrobacterium produced the highest myoglobin accumulation, with individual seed stocks ranging from 0.32% to 1.57% TSP. Several independent events consistently exceeded 0.8%, and the top-performing line reached 1.57 ± 0.63%, representing a ∼50-fold increase relative to WT controls. This indicates that pRG13 was the most efficient plant transformation vector for driving high-level myoglobin expression in pea seeds. The pRG14 (PPHAS Promoter) construct produced intermediate expression levels, with accumulation spanning 0.13% to 0.83% TSP. Although lower than pRG13 on average, several lines exceeded 0.6%, suggesting that pRG14 can achieve moderate expression but is more variable between events. In contrast, pRG15 (CaMV35S-TEV Promoter)-transformed lines accumulated only 0.09–0.14% TSP, which is near background levels. This indicates that pRG15 is substantially less effective at supporting myoglobin expression compared to pRG13 and pRG14, possibly due to differences in promoter strength, codon usage, or construct stability.

Finally, seeds obtained from events expressing pRG14 + pRG16 via particle bombardment exhibited a broad distribution of expression levels (0.03–0.77% TSP). Several lines reached levels comparable to the top-performing Agrobacterium-derived pRG14 lines, but others expressed at very low or near-background levels. This variability suggests that biolistic delivery can enable relatively high expression, but integration site effects and transgene stability may lead to inconsistent outcomes across independent events.

Overall, these findings demonstrate that construct design strongly influences myoglobin accumulation in pea seeds, with 7S globulin promoters fused with the Tobacco Etch virus translational enhancer providing the most robust expression. Transformation method also contributed to variability, with Agrobacterium-mediated delivery producing more consistently high-expressing lines compared to bombardment. Segregation ratios, transgene copy number, and potential chimerism were not systematically evaluated in this study. Given previous reports of non-Mendelian segregation and mosaic integration following embryonic-axis transformation in pea, these factors may contribute to the observed variability in expression.

These results establish a foundation for further optimization of seed-based myoglobin production systems, with implications for both functional food biotechnology and plant-based alternative protein development.

## Conclusions

This study demonstrates stable seed-specific expression of myoglobin in *Pisum sativum* using both Agrobacterium-mediated transformation and particle bombardment. Among the regulatory elements tested, myoglobin expressed under the regulation of 7S promoter fused with TEV translational enhancer consistently yielded the highest levels of myoglobin accumulation, with individual lines reaching up to 1.57% of total soluble protein. These results reinforce the role of *Pisum sativum* as a promising host for molecular farming of functional animal proteins. To the best of our knowledge, the expression of the heme-containing myoglobin protein represents the first study for producing a potential food product in pea seeds. This is an important development because pea processing and extrusion for use as food ingredients is already a mature industry (Shanthakumar et al. 2022). Moreover, pea protein augmented with myoglobin is expected to enhance the nutritional profile. Future efforts aimed at promoter selection, codon optimization, and marker-free selection systems may further enhance yield and stability, paving the way for the development of plant-based platforms to supply functional heme proteins for food, nutrition, and biotechnological applications.

## Statements and Declarations

### Competing Interests

AD served as the Chief Science Officer at Moolec from 10/01/2020 to 07/31/2025 with active conflict of interest disclosures on file at Washington State University and Texas A&M University. The funder provided the design of the plant transformation vectors and BWB and NG assessed the myoglobin protein content in confirmed transgenic lines which for which they have been appropriately credited as co-authors. Besides that, Moolec Science Limited had no role in experimental design, data collection and analysis.

### Author Contributions

AD and RG designed the study. RG, KA, NT, KC, DV, LW, RH, JH performed experiments and generated data. BWB and NG at Moolec Science Limited performed experiments to generate myoglobin protein expression data. RG, BWB and AD developed the first draft. AD finalized the manuscript. All authors read and approved of the final manuscript.

### Funding

This study was supported by Moolec Science Limited through a Sponsored Research Project Agreement ORSO #140369-00 to AD while he was at Washington State University (WSU), entered into by and between Washington State University (WSU) and Moolec Science Limited, dated January 29, 2021, as amended. The work was also supported in part by Texas A&M AgriLife Hatch Project #TEX0-9950-0 and startup funds from Texas A&M AgriLife Research and Texas A&M University to AD.

## Acknowledgements

Not applicable.

## References

Aragão FJL, Barros LMG, Brasileiro ACM, et al (1996) Inheritance of foreign genes in transgenic bean (Phaseolus vulgaris L.) co-transformed via particle bombardment. Theor Appl Genet 93:142–150. 10.1007/BF00225739

Bean SJ, Gooding PS, Mullineaux PM, Davies DR (1997) A simple system for pea transformation. Plant Cell Rep 16:513–519. 10.1007/BF01142315

Carlsson MLR, Kanagarajan S, Bülow L, Zhu L-H (2020) Plant based production of myoglobin - a novel source of the muscle heme-protein. Sci Rep 10:920. 10.1038/s41598-020-57565-y

Chen H, Osuna D, Colville L, et al (2013) Transcriptome-Wide Mapping of Pea Seed Ageing Reveals a Pivotal Role for Genes Related to Oxidative Stress and Programmed Cell Death. PLoS One 8:e78471. 10.1371/JOURNAL.PONE.0078471

Christou P (1990) Soybean transformation by electric discharge particle acceleration. Physiol Plant 79:210–212. 10.1111/J.1399-3054.1990.TB05889.X;CTYPE:STRING:JOURNAL

Davis B (2025) The Power of Pea Protein: Why It’s Leading the Charge in 2025 - Plant Based World Pulse. https://plantbasedworldpulse.com/the-power-of-pea-protein-why-its-leading-the-charge-in-2025/. Accessed 9 Nov 2025

Day L, Cakebread JA, Loveday SM (2022) Food proteins from animals and plants: Differences in the nutritional and functional properties. Trends Food Sci Technol 119:428–442. 10.1016/J.TIFS.2021.12.020

Devaere J, De Winne A, Dewulf L, et al (2022) Improving the aromatic profile of plant-based meat alternatives: effect of myoglobin addition on volatiles. Foods 11:1985

Dhingra A, Daniell H (2006) Chloroplast genetic engineering via organogenesis or somatic embryogenesis. Methods Mol Biol 323:245–262. 10.1385/1-59745-003-0:245

Fan Y, Li W, Wang J, et al (2011) Efficient production of human acidic fibroblast growth factor in pea (Pisum sativum L.) plants by agroinfection of germinated seeds. BMC Biotechnology 2011 11:1 11:1–9. 10.1186/1472-6750-11-45

Fauteux F, Strömvik M V (2009) Seed storage protein gene promoters contain conserved DNA motifs in Brassicaceae, Fabaceae and Poaceae. BMC Plant Biol 9:126. 10.1186/1471-2229-9-126

Fraeye I, Kratka M, Vandenburgh H, Thorrez L (2020) Sensorial and nutritional aspects of cultured meat in comparison to traditional meat: much to be inferred. Front Nutr 7:35

Grant JE, Cooper PA, McAra AE, Frew TJ (1995) Transformation of peas (Pisum sativum L.) using immature cotyledons. Plant Cell Reports 1995 15:3 15:254–258. 10.1007/BF00193730

Henchion M, Hayes M, Mullen AM, et al (2017) Future Protein Supply and Demand: Strategies and Factors Influencing a Sustainable Equilibrium. Foods 2017, Vol 6, Page 53 6:53. 10.3390/FOODS6070053

Indurker S, Misra HS, Eapen S (2007) Genetic transformation of chickpea (Cicer arietinum L.) with insecticidal crystal protein gene using particle gun bombardment. Plant Cell Reports 2006 26:6 26:755–763. 10.1007/S00299-006-0283-6

Jackson MA, Anderson DJ, Birch RG (2012) Comparison of Agrobacterium and particle bombardment using whole plasmid or minimal cassette for production of high-expressing, low-copy transgenic plants. Transgenic Research 2012 22:1 22:143–151. 10.1007/S11248-012-9639-6

Jiwan D, Roalson EH, Main D, Dhingra A (2013) Antisense expression of peach mildew resistance locus O (PpMlo1) gene confers cross-species resistance to powdery mildew in Fragaria x ananassa. Transgenic Res 22:1119–1131

Jordan MC, Hobbs SLA (1993) Evaluation of a cotyledonary node regeneration system forAgrobacterium-mediated transformation of pea (Pisum sativum L.). In Vitro Cellular & Developmental Biology - Plant 1993 29:2 29:77–82. 10.1007/BF02632256

Kaur A, Sharma M, Sharma C, et al (2017) Genetic Transformation Of Pigeonpea Through Particle Gun And Agrobacterium Using Cry 1 ac Transgene. Plant Cell Tissue Organ Cult 127:717–727. 10.1007/S11240-016-1055-9

Kaur R, Donoso T, Scheske C, et al (2022a) Highly Efficient and Reproducible Genetic Transformation in Pea for Targeted Trait Improvement. ACS Agricultural Science & Technology 2:780–787. 10.1021/acsagscitech.2c00084

Kaur R, Donoso T, Scheske C, et al (2022b) Highly Efficient and Reproducible Genetic Transformation in Pea for Targeted Trait Improvement. ACS Agricultural Science & Technology 2:780–787. 10.1021/ACSAGSCITECH.2C00084

Li G, Liu R, Xu R, et al (2023) Development of an Agrobacterium-mediated CRISPR/Cas9 system in pea (Pisum sativum L.). Crop J 11:132–139. 10.1016/j.cj.2022.04.011

Liu N, Zhang G, Xu S, et al (2015) Comparative transcriptomic analyses of vegetable and grain pea (Pisum sativum L.) seed development. Front Plant Sci 6:1–14. 10.3389/FPLS.2015.01039/BIBTEX

Ludvíková M, Griga M (2022) Pea transformation: History, current status and challenges. Czech Journal of Genetics and Plant Breeding 58:127–161. 10.17221/24/2022-CJGPB

Malovichko Y V., Shtark OY, Vasileva EN, et al (2020) Transcriptomic Insights into Mechanisms of Early Seed Maturation in the Garden Pea (Pisum sativum L.). Cells 2020, Vol 9, Page 779 9:779. 10.3390/CELLS9030779

Messina M, Messina V (2024) Health and functional advantages of cheese containing soy protein and soybean-derived casein. Front Plant Sci 15:1407506. 10.3389/FPLS.2024.1407506/FULL

Mikschofsky H, Schirrmeier H, Keil GM, et al (2009) Pea-derived vaccines demonstrate high immunogenicity and protection in rabbits against rabbit haemorrhagic disease virus. Plant Biotechnol J 7:537–549. 10.1111/J.1467-7652.2009.00422.X

Nadolska-Orczyk A, Orczyk W (2000) Study of the factors influencing Agrobacterium-mediated transformation of pea (Pisum sativum L.). Molecular Breeding 6:185–194. 10.1023/A:1009679908948/METRICS

Nandi S, Suzuki YA, Huang J, et al (2002) Expression of human lactoferrin in transgenic rice grains for the application in infant formula. Plant Science 163:713–722. 10.1016/S0168-9452(02)00165-6

Paladini G, Salinas M, Dhingra A, et al (2024) US20240352475A1 - High expression of animal heme protein in plants - Google Patents. https://patents.google.com/patent/US20240352475A1/en

Partanen M, Liikonen V, Väkeväinen K, et al (2025) Digestion, Metabolism, and Health Effects of Plant Proteins and Their Food Formulations: A Systematic Scoping Review of Clinical Postprandial Studies and in vitro Methods. Food Reviews International. 10.1080/87559129.2025.2525430;CTYPE:STRING:JOURNAL

Perrin Y, Vaquero C, Gerrard I, et al (2000) Transgenic pea seeds as bioreactors for the production of a single-chain Fv fragment (scFv) antibody used in cancer diagnosis and therapy. Molecular Breeding 6:345–352. 10.1023/A:1009657701588/METRICS

Philip R, Darnowski DW, Maughan PJ, Vodkin LO (2001) Processing and localization of bovine β-casein expressed in transgenic soybean seeds under control of a soybean lectin expression cassette. Plant Science 161:323–335. 10.1016/S0168-9452(01)00420-4

Pniewski T, Kapusta J (2005) Efficiency of transformation of Polish cultivars of pea (Pisum sativum L.) with various regeneration capacity by using hypervirulent Agrobacterium tumefaciens strains. J Appl Genet 46:139–47

Saalbach I, Giersberg M, Conrad UDO (2001) High-level expression of a single-chain Fv fragment (scFv) antibody in transgenic pea seeds. J Plant Physiol 158:529–533. 10.1078/0176-1617-00366

Schroeder HE, Schotz AH, Wardley-Richardson T, et al (1993) Transformation and Regeneration of Two Cultivars of Pea (Pisum sativum L.). Plant Physiol 101:751–757. 10.1104/pp.101.3.751

Shanthakumar P, Klepacka J, Bains A, et al (2022) The Current Situation of Pea Protein and Its Application in the Food Industry. Molecules 27. 10.3390/MOLECULES27165354

Shoseyov O, Posen Y, Grynspan F (2013) Human recombinant type i collagen produced in plants. Tissue Eng Part A 19:1527–1533. 10.1089/TEN.TEA.2012.0347

Soh BXP, Smith NW, Von Hurst PR, McNabb WC (2025) Achieving High Protein Quality Is a Challenge in Vegan Diets: A Narrative Review. Nutr Rev 83:e2063–e2081. 10.1093/NUTRIT/NUAE176

Suman SP, Joseph P (2013) Myoglobin chemistry and meat color. Annu Rev Food Sci Technol 4:79–99

Tusé D, McNulty M, McDonald KA, Buchman LW (2024) A review and outlook on expression of animal proteins in plants. Front Plant Sci 15:1426239. 10.3389/FPLS.2024.1426239/FULL

Vianna GR, Aragão FJ, Rech EL (2011) A minimal DNA cassette as a vector for genetic transformation of soybean (Glycine max). Genet Mol Res 10:382–390. 10.4238/VOL10-1GMR1058

Wandelt C, Knibb W, Schroeder HE, et al (1991) The Expression of an Ovalbumin and a Seed Protein Gene in the Leaves of Transgenic Plants. Plant Molecular Biology 2 471–478. 10.1007/978-1-4615-3304-7_45

Williamson-Benavides BA, Parida A, Manning J, et al (2025) Efficient Stable Genetic Transformation of Pea (Pisum sativum). 10.21203/RS.3.RS-7841977/V1

Williamson-Benavides BA, Sharpe RM, Nelson G, et al (2020) Identification of Fusarium solani f. sp. pisi (Fsp) Responsive Genes in Pisum sativum. Front Genet 11:. 10.3389/FGENE.2020.00950

Xu H, Guo Y, Qiu L, Ran Y (2022) Progress in Soybean Genetic Transformation Over the Last Decade. Front Plant Sci 13:900318. 10.3389/FPLS.2022.900318/FULL

Yang T, Liu R, Luo Y, et al (2022) Improved pea reference genome and pan-genome highlight genomic features and evolutionary characteristics. Nat Genet 54:1553–1563. 10.1038/s41588-022-01172-2

Zakharov A, Giersberg M, Hosein F, et al (2004) Seed-specific promoters direct gene expression in non-seed tissue. J Exp Bot 55:1463–1471. 10.1093/jxb/erh158

Zimmermann J, Saalbach I, Jahn D, et al (2009) Antibody expressing pea seeds as fodder for prevention of gastrointestinal parasitic infections in chickens. BMC Biotechnol 9:79. 10.1186/1472-6750-9-79

